# Evolutionary dynamics of neoantigens in growing tumours

**DOI:** 10.1101/536433

**Authors:** Eszter Lakatos, Marc J. Williams, Ryan O. Schenck, William C. H. Cross, Jacob Househam, Benjamin Werner, Chandler Gatenbee, Mark Robertson-Tessi, Chris P. Barnes, Alexander R. A. Anderson, Andrea Sottoriva, Trevor A. Graham

## Abstract

Cancer evolution is driven by the acquisition of somatic mutations that provide cells with a beneficial phenotype in a changing microenvironment. However, mutations that give rise to neoantigens, novel cancer–specific peptides that elicit an immune response, are likely to be disadvantageous. Here we show how the clonal structure and immunogenotype of growing tumours is shaped by negative selection in response to neoantigenic mutations. We construct a mathematical model of neoantigen evolution in a growing tumour, and verify the model using genomic sequencing data. The model predicts that, in the absence of active immune escape mechanisms, tumours either evolve clonal neoantigens (antigen– ‘hot’), or have no clonally– expanded neoantigens at all (antigen– ‘cold’), whereas antigen– ‘warm’ tumours (with high frequency subclonal neoantigens) form only following the evolution of immune evasion. Counterintuitively, strong negative selection for neoantigens during tumour formation leads to an increased number of antigen– warm or – hot tumours, as a consequence of selective pressure for immune escape. Further, we show that the clone size distribution under negative selection is effectively– neutral, and moreover, that stronger negative selection paradoxically leads to more neutral– like dynamics. Analysis of antigen clone sizes and immune escape in colorectal cancer exome sequencing data confirms these results. Overall, we provide and verify a mathematical framework to understand the evolutionary dynamics and clonality of neoantigens in human cancers that may inform patient– specific immunotherapy decision– making.

## INTRODUCTION

Mutations accrue throughout tumour development and provide ‘fuel for the fire’ of cancer evolution. *Driver* mutations cause evolutionary adaptive (beneficial) changes to the phenotype of the cancer cell within its microenvironment. However, mutations can also hinder tumour evolution if they lead to an anti– tumour immune response, via the generation of neoantigens, novel peptides presented on the cell’s surface recognised as ‘non– self’ by tumour– infiltrating lymphocytes of the adaptive immune system^1,2^. Highlighting the broad importance of the immune system in tumour evolution is the observation that the abundance of immune– infiltration in tumours correlates with patient survival^3^, and with the success of immunotherapy approaches that lead to an anti– tumour immune response^4^.

The landscape of neoantigen– associated mutations is shaped by ecological and evolutionary interactions between a tumour and its microenvironment^1,5^. In a scenario where the tumour is isolated from the immune system, neoantigens accumulate as a ‘side– effect’ of mutation acquisition^6^, and are expected to follow neutral evolutionary dynamics^7^. Consequently, by default the overall mutational load of a tumour is expected to contain a proportion of neoantigens that will elicit an immune response. The ultimate consequence of the immune response is to kill the antigenic cells, such that only the non– antigenic cells survive – a process often referred to as *immuno–editing*^1^. In face of this selective pressure, tumour cells benefit if they evolve a mechanism to inhibit the immune system’s ability to recognise or react to cancer– associated antigens: these mechanisms are termed *immune escape* or *immune evasion* mechanisms^6,8,9^. Together this means that the neoantigenic repertoire found in tumours has been evolutionary– selected to survive in the tumour– specific immune microenvironment^10^. Adaptive evolution to immune control is recognised as a ‘hallmark of cancer’^11^.

Therapies that counteract immune escape by targeting the (re)activation of immune response can achieve an exceptional success in cancers^12,13^, especially in those with a high mutational load^14–16^. For example, pharmacological interference with inhibitory immune checkpoint such as CTLA– 4 or PD– L1 restores cytotoxicity of existing cancer– specific T– cells and hence initiates a potent response against cancer– associated neoantigens, often achieving long– term remission even in advanced cancers^2,17,18^. However, accurate prediction of immune– therapy responsiveness remains challenging, as a significant number of patients do not experience a complete or even partial response, regardless of a high mutational load and the presence of molecular markers of immune escape^19,20^. Therefore, understanding the evolutionary processes that contributed to an individual cancer’s neoantigen landscape may usefully inform therapeutic decision–making.

The evolutionary dynamics of tumour development are encoded in the pattern of intra– tumour genetic heterogeneity. Specifically, strongly beneficial or strongly disadvantageous mutations by definition affect the net reproduction of cells that carry them, and consequently will be present in, respectively, higher or lower numbers of cells in the final population. Utilising this idea, mathematical modelling of the clone size distribution of mutations in tumours (as measured by the distribution of variant allele frequencies (VAF)) facilitates inference of the evolutionary dynamics that shaped the mutational landscape of individual tumours^21^. We have previously developed models to infer the presence of strong positive selection or effectively– neutral dynamics from sequencing data^7,22^ and theoretical analysis of selection have been extended for disadvantageous mutation sites^23,24^. However, the influence of negative selection acting on neoantigens, as observed in the clone size distribution of cancers, remains to be determined^25^.

Here we used a stochastic modelling approach to study how the clonal structure and immunological phenotype of growing tumours is shaped by negative selection in response to neoantigenic mutations. We establish the characteristic clone size distribution under pervasive negative selection for neoantigens, and determine how this is changed by immune– escape. We use colorectal cancer sequencing data to verify these predictions.

## RESULTS

### Mathematical model of tumour growth and mutation accumulation

We created a stochastic branching process based mathematical model of negatively– selected neoantigen evolution during tumour growth (Figure 1A). In the model, tumour evolution was initiated by a single transformed cell that produced two surviving offspring at birth rate *b* per unit time, and for simplicity, we set *b*=1. Cells in clone *i* died at rate *d_i_* per unit time, where the death rate increased with the neoantigen burden of the clone. Each time a cell divided, it acquired new unique mutations at overall rate *µ* (Poisson distribution), which were assigned as neoantigens at rate *p*_*na*_, or as passengers (evolutionary neutral) at rate 1– *p*_*na*_. Neoantigens caused the death rate *d*_*i*_ of the lineage to increase from a basal rate of *d_b_*=0.1 to a higher value determined by the strength of negative selection against each new neoantigen s, and the number of neoantigens harboured in the lineage *n_i_*. We defined the selective (dis)advantage of a subclone by its effective proliferation rate (the difference of its birth and death rate), as compared to a non– immunogenic clone:

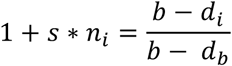

**Figure 1:**
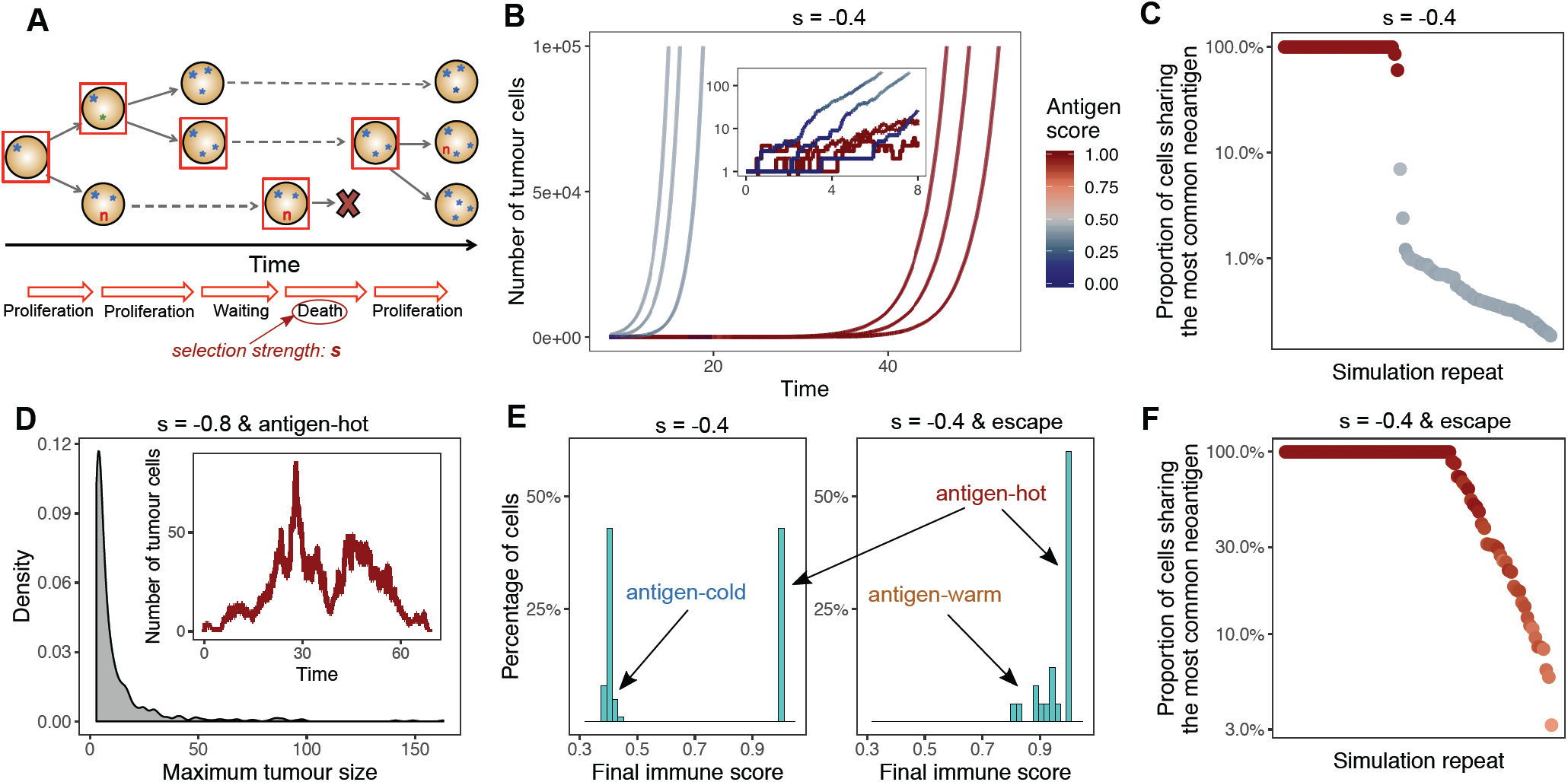
Tumour growth model predicts two distinct types of immune phenotypes and the necessity of immune escape. **(A)** Schematic representation of a simulation of the tumour growth model. Filled circles represent cells, and characters represent (sets of) mutations harboured in each cell: blue stars are neutral, red ‘**n**’s are neoantigen-associated mutations. The horizontal axis indicates time. Starting from a single progenitor cell, in each step a tumour cell is randomly selected (denoted by the red square) and undergoes proliferation or death. Daughter cells inherit the mutations of the parent cell and also accumulate new alterations. The probability of death events depends on the selection pressure, s. **(B)** Growth curve of six simulated tumours. The colour of the line shows the antigenicity of the tumour population at that time. The inset depicts the first segment of the growth curves on a logarithmic scale. **(C)** The proportion of tumour cells sharing the highest frequency antigenic mutation in 100 tumours simulated at selection strength s=-0.4, each dot representing a separate tumour. The colour of the dots shows the final antigenicity of the tumour. **(D)** Distribution of the maximum tumour size (measured as the highest number of tumour cells) reached by simulated antigen hot tumours under strong negative selection (s=-0.8). The inset depicts the growth and extinction of such a tumour. **(E)** Distribution of final antigenicity values measured over 100 identical tumours at selection s=-0.4, without (left) and with (right) clonal immune escape. **(F)** Frequency of the most shared antigenic mutation in 100 immune escaped tumours at selection pressure s=-0.4. The colour of the dots shows the final antigenicity of the tumour (scale in panel B).

where *s*=0 for no selection pressure (neoantigens carry no disadvantage, neutral evolution), and *s*<0 for selection against neoantigens (following ^23^). Consequently, the death rate of a subclone was computed as:

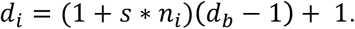

This neoantigen number– dependent increase in the clone death rate represented an aggregate of the many factors that lead to the negative selection of neoantigens, including; (i) sufficient presentation of neoantigens on the cell surface; (ii) recognition of neoantigens by T– cell; (iii) T– cell killing efficiency. Different values of selection strength, s, included in the death rate should be interpreted as a measure of the efficacy of immune predation.

Tumour growth was simulated until the tumour reached a predefined population size (approximating a clinically detectable size) or until a sufficiently long time elapsed without tumour establishment (corresponding to no cancer formation in a patient’s lifetime). We examined the clonal structure of the tumour, both of neutral and neoantigenic mutations, throughout the simulation of tumour growth.

### Modelling predicts tumours are distinctively antigen–cold or hot

We simulated neoantigen evolution during tumour growth and examined the neoantigen burden of the resulting tumours. To quantify population– level immunogenicity, we defined the ‘antigen score’ of a tumour as the proportion of tumour cells that were antigenic (carried at least one neoantigen– associated mutation). Large variations in antigenicity were observed between tumours simulated under identical conditions (e.g. constant antigen production rate *p*_*na*_=0.1, and negative selection strength s=– 0.4) (Figure 1B). Specifically, the simulated tumours divided into two clear groups: ‘antigen– hot’ and ‘antigen– cold’. Antigen– hot tumours had an antigen score of 1, corresponding to every tumour cell carrying at least one antigenic mutation, whereas in antigen– cold tumours the majority of cells lacked immunogenic mutations. We note that, as new neoantigens are generated at a constant rate, there was always a non– zero population of cells (multiple small subclones) that carried neoantigens. The proportion of antigen– hot tumours depended on the neoantigen– production rate and selection strength (Figure S1A): decreased production rate and increased negative selection for neoantigens diminished the probability of observing an antigen– hot tumour. In antigen– cold tumours, the proportion of neoantigen– carrying cells similarly decreased with production rate and increased negative selection against antigens.

In antigen– hot tumours the most common antigenic mutation was always clonal or very close to complete clonality (Figure 1C), showing that these tumours arose if, through random drift, a neoantigen– carrier cell was ‘lucky’ and became the progenitor of the final tumour. Within an antigen– hot tumour, the entire tumour cell population is then subject to the increased death rate (due to immune predation) resulting from the clonal neoantigen, and correspondingly, antigen–hot tumours had a slower net growth rate and so required a longer time to reach detectable size. In antigen– cold tumours, the immunogenic population was composed of a highly heterogeneous collection of small subclones wherein no individual neoantigen could be found in more than 10% of cells, despite over 40% of the total tumour cells carrying a neoantigen (Figure 1C). In these cases, a non– antigenic clone founded the tumour, and later arising neoantigens experienced immuno– editing preventing their clonal expansion. Non– antigenic clones experienced no negative selection, and so expanded unperturbed. Therefore, due to this disparity in growth between immunogenic and non–immunogenic parts of the tumour, the clones carrying neoantigens declined in frequency over time.

We explored the overall antigen burden under low, medium and high negative selection (s=– 0.1,– 0.4 and – 0.8, respectively). We simulated error– prone sequencing (see Methods), and measured the burden of each simulated tumour as the number of antigenic mutations thus detected. The strength of the selection pressure led to highly variable detectable antigen burden distributions (Figure S1B), with a negative correlation between the average antigen burden and selection strength, as expected.

We observed that strong negative selection for neoantigens led to the evolution of only antigen– cold tumours (Figure S1A), implying that tumours carrying clonal antigens are not viable when immune selection is strong. To confirm this, we simulated antigen– hot tumours, by introducing an antigenic mutation in the founder cell of the population, and measured resulting tumour growth under high negative selection pressure (s=– 0.8). In this selective regime, the acquisition of a second antigen lead to the rapid eradication of the double– antigen clone; consequently, tumour size was severely restricted (Figure 1D). We observed the same trend when simulating hyper– mutated tumours that generated 4– 10 exonic mutations per cell division. The high rate of neoantigen production under this scenario led to the number of non– antigenic cells in a given clone declining over cell divisions, and growing subclones rapidly becoming neoantigen– hot and thus leading to tumour death via immune predation (Figure S2A), recapitulating the behaviour proposed previously^26^.

### Modelling predicts immune escape leads to increased antigen– hot and – warm tumours

We next investigated the evolutionary consequence of immune escape. We simulated the scenario where an *immune escape alteration* in one cell (potentially the founding cell of the tumour) renders that cell and its descendants less susceptible to immune predation. Known immune escape mechanisms include mutations to antigen presenting machinery and expression of immune checkpoint molecules^27,28^. While some immune evasion mechanisms are considered phenotypic, we argued that they still can partially originate from underlying heritable changes (e.g. copy number alteration of the PD– L1 gene), and hence can be modelled in our mutational framework. We set the death rate of immune escaped cells to the baseline non– immunogenic death rate, *d*_*b*_, irrespective of the antigenic mutations harboured by the cell. Therefore, immune escape effectively meant the clone returned to evolve under neutral dynamics.

If the founder cell of the tumour contained an escape mutation (*clonal immune escape*) there was no longer separation into antigen– hot or – cold tumours (Figure 1E and S1D). Stochastic evolution of immune escape could ‘rescue’ a fledging tumour from extinction and lead to a cancer of detectable size even under conditions previously found to be unviable (Figure S2). Furthermore, the resulting tumours were often either antigen– hot or antigen– warm (containing high frequency neoantigens), by virtue of the majority of cells in the tumour carrying (multiple) neoantigens, irrespective of selection strength (Figure 1E & F and S1C & E). Both hot and warm tumours appeared more frequently than for simulations where immune– escape was not permitted.

Taken together, these results predicted that tumours that grow to detectable size in an environment with high levels of immune surveillance are likely to be enriched for immune escape, and consequently show an unexpectedly high proportion of antigen–warm and –hot tumours. These observations will be more pronounced in hyper–mutated samples where the selective pressure to acquire immune escape will be stronger due to the continual acquisition of neoantigens.

### Most colorectal cancers are antigen– hot and enriched for immune evasion

To verify the mathematical model, we explored the clonality of neoantigens in colorectal cancer (CRC, combined colon and rectal adenocarcinoma dataset) samples from The Cancer Genome Atlas (TCGA) database. We focused on CRC because of the prevalence of mutator phenotypes: cancers with polymerase– ε mutation (POLE – very high mutation rate), with microsatellite instability (MSI – high mutation rate), and microsatellite stable tumours (MSS – normal mutation rate). Therefore, CRC provides a good model to explore the effect of different tumour– immune environments. Furthermore, therapy based on immune checkpoint inhibitors has been successfully applied for MSI tumours, but the efficacy is still not known for the majority of CRC cases^29^.

CRC samples filtered for high sequencing depth and purity were first HLA– typed in silico from associated blood– derived normals^30^, and neoantigens called from tumours using the NeoPredPipe pipeline^31^. We filtered neoantigen calls based on predicted HLA binding affinity and similarity to known antigens to leave only neoantigens predicted to be highly antigenic^32^. We then assessed the cancer cell fraction (CCF) of these ‘strong’ neoantigens (see Methods).

The vast majority of tumours (90%) had clonal neoantigens and so were defined as “antigen– hot” (Figure 2A). There was a more than two– fold order of magnitude difference in the neoantigen burden between tumours (Figure 2B) and the level of T– cell presence (as measured from paired RNA– seq data ^33^) – a proxy measure of the intensity of immune surveillance, represented in our model by the negative selection strength s – varied dramatically between samples. Colorectal cancers with low or medium T– cell infiltration (putative small or moderate s) tended to have proportionally fewer clonal neoantigens than tumours with high levels of T–cells infiltrate (putative high s) (Figure 2B), suggesting a critical role of immune escape in the initial evolution of these tumours. These findings are in accordance with previous studies exploring the connection between neoantigen burden and the efficacy of therapies targeting immune–escape mechanisms^34^.

**Figure 2:**
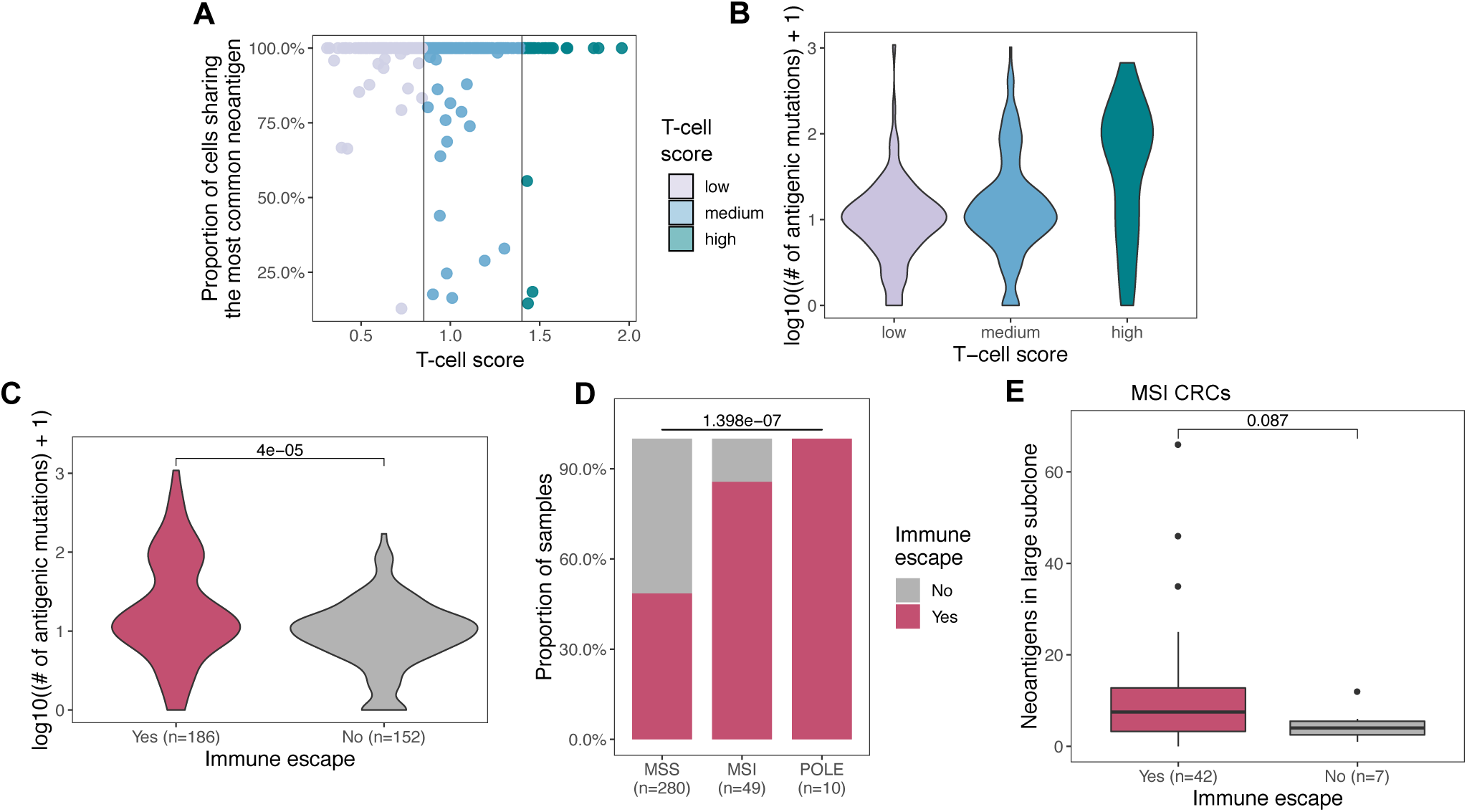
Colorectal tumours from TCGA are antigen-hot and enriched for immune escape. **(A)** The proportion of tumour cells sharing the most common neoantigen-associated mutation in each tumour in the TCGA CRC domain, each dot representing a single tumour. Tumours are ordered according to their T-cell score, and divided up to low (light purple), medium (blue) and high (green) immune infiltration categories. **(B)** Distribution of the number of detected neoantigen-associated mutations TCGA CRCs with low, medium and high immune infiltration (T-cell average) score. **(C)** Distribution of the number of detected antigenic mutations in TCGA samples with and without detected immune escape (magenta and grey violins, respectively). The p-value of a Mann-Whitney test comparing the means of the two populations is reported above the violin plots. **(D)** Prevalence of immune escape in the colorectal TCGA domain divided into MSS, MSI and POLE samples. Chi-squared test is indicated on top of the panel. **(E)** Number of neoantigens present in high subclonal proportions (between 30% and 60% of cells) in MSI TCGA CRC samples with and without immune escape. The p-value of one-sided Mann-Whitney test is reported above the plot.

We then sought evidence of immune escape in these cancers. Using exome and RNAseq data, we tested for the presence of three types of immune escape mechanisms: (i) somatic mutations in either one of the HLA alleles or in the B2M gene^30,33^; (ii) loss of an HLA haplotype through loss of heterozygosity (LOH) in the corresponding genomic locus^28^; and (iii) PD– L1 or CTLA– 4 over–expression^35^. Confirming the prediction of the simulation, tumours with immune–escape had a higher antigen burden, and the majority of highly antigenic tumours (with antigen burden >100) were immune– escaped (Figure 2C).

We next explored the prevalence of immune escape as a function of the underlying mutation rate, and consequently, neoantigen burden, by comparing the frequency of escape mechanisms detected in cancers with POLE, MSI, and MSS genotype. Overall, 55% of all cancers showed evidence of at least one escape mechanism, with increased prevalence of escape in MSI (86%) and POLE (100%) cases. We note that the extent of the impact of these escape alterations is not always known – especially for non– synonymous mutations affecting HLA – but we argued that they might influence the evolution of the antigen landscape. We found that hyper– mutated tumours also showed significantly different patterns of immune escape (Figure 2D and Figure S3), in agreement with previous studies^16,33^. They were dominated by immune checkpoint over–expression and point mutations in important immune genes, compared to an enrichment of chromosomal loss of HLA in MSS tumours, in agreement with the high percentage of chromosomal aberrations observed in MSS CRC^36^. Further work is needed to determine whether these differences between mutational subtypes arose solely as a result of underlying mutational frequency, or different selective pressures on immune evasion.

We also tested if the presence of immune escape had altered the strength of immuno– editing. The interplay between negative selection and immune evasion had a most profound effect in the regime of large subclones (Figure 1C & F): neoantigen– associated mutations are most significantly under– represented amongst subclonal mutations at high CCF. Therefore, we compared the number of neoantigens at high CCF (present in 30% to 60% of cells) between MSI CRC cases with and without immune escape. High–CCF subclonal neoantigens were enriched in hyper– mutated cases with immune escape (Figure 2E), consistent with immune–surveillance and immuno– editing strongly shaping the clonal structure of hyper– mutated tumours.

Together, these data suggest that colorectal cancers usually evolve in the face of stringent immune– selective pressures (strong immuno– editing) and consequently that immune–escape is frequently selected for at the onset of tumour growth, allowing for tumours with high antigen load.

### Subclonal immune escape shapes local neoantigen evolution and response to therapy

Next, we explored the evidence for subclonal immune escape in our previously published multi– region sequenced colorectal tumour dataset^37^. Overall, loss of heterozygosity (LOH) at HLA loci was found in 5/10 (50%) carcinomas and 1/6 (17%) adenomas and no HLA or B2M mutations were detected (immune checkpoint over–expression could not be inferred as only genomic data was available) (Figure 3A). In four of the cases with evidence of immune escape, we detected HLA LOH events that were present only subclonally, in spatially distinct region(s) of the tumour (Figure 3B).

**Figure 3:**
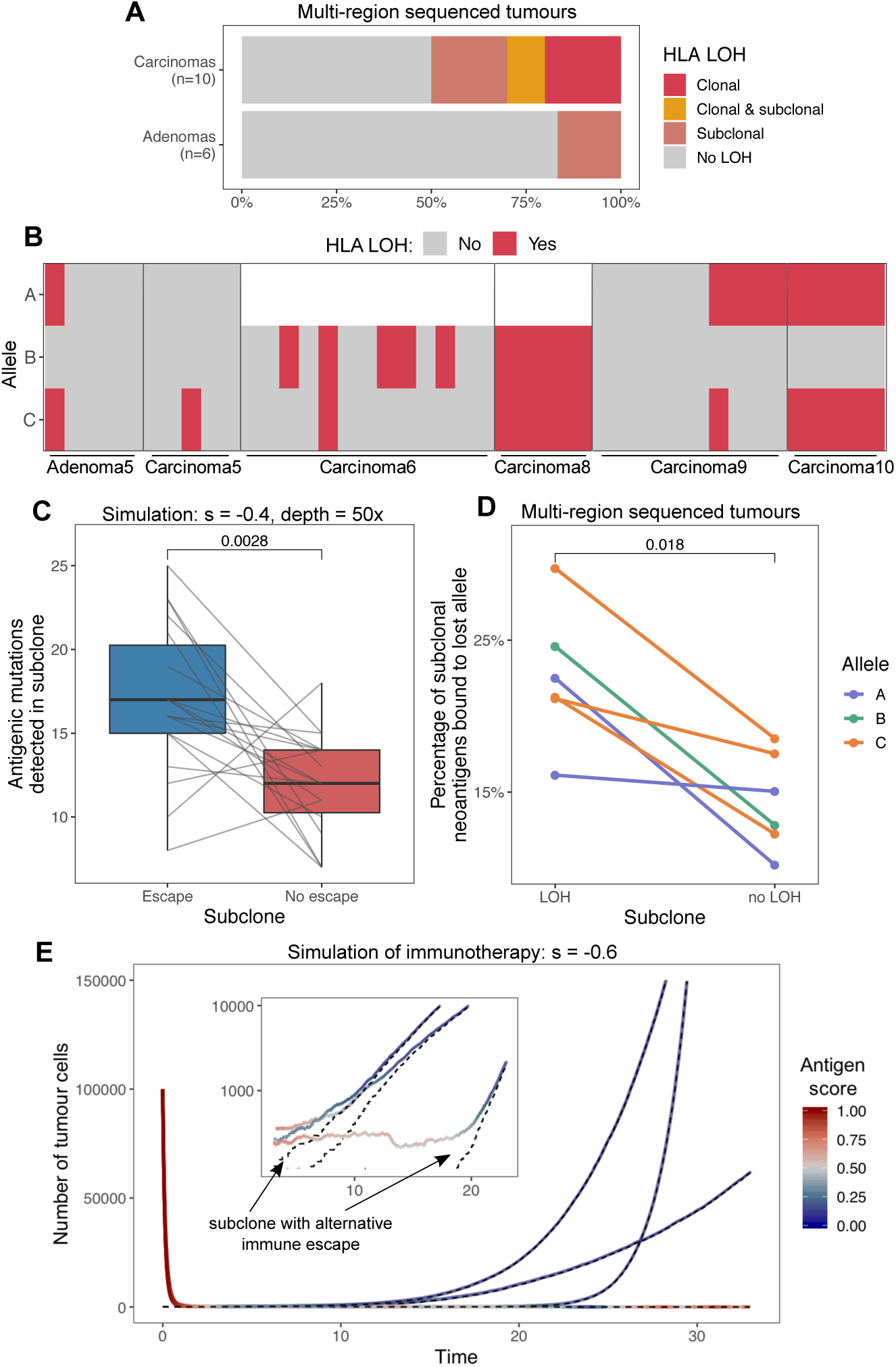
Subclonal immune evasion shapes neoantigen landscape and tumour growth after therapy. **(A)** Immune escape through LOH at an HLA locus in the multi-region sequenced colorectal cohort. LOH events are divided up according to whether the alteration is detected in all (clonal) or not all (subclonal) biopsies. **(B)** HLA LOH in individual biopsies in tumours with at least one subclonal or clonal loss event. Unfilled boxes represent homozygous HLA alleles. **(C)** The number of antigenic mutations detected in two distinct (with and without immune escape) subclones of a simulated tumour. Antigenic mutations are detected after simulated sequencing with read depth of 50x. **(D)** The proportion of all neoantigens binding to the HLA allele lost in the LOH event in the colorectal tumours that show subclonal HLA LOH. P-values of one-sided Wilcoxon signed-rank tests are reported on (C) and (D). **(E)** Growth curve of simulated tumours following anti-PD-L1-type immunotherapy. The tumours have previously developed active immune evasion, but also harbour a small subclone with different escape mechanism. Black dashed lines show the number of cells in this subclone over time. The inset shows growth around the point when the subclone takes over, on a logarithmic scale.

Simulations of subclonal immune– evasion in our model predicted that subclones should become proportionally enriched for neoantigens following immune evasion (Figure 3C), with the detectability of this effect dependent on the sequencing depth and strength of negative selection for neoantigens (Methods). To test this prediction, we compared neoantigen burdens in immune– escaped versus not– escaped tumour subregions to explore the effects of spatial heterogeneity in immune selective pressures within individual tumours. We hypothesised that in case of HLA– LOH, antigens binding to the (subclonally) lost HLA allele would similarly experience a release of selection pressure and rise to higher frequencies. Consistent with this hypothesis, a significantly higher proportion of detected neoantigens were predicted to bind to the lost allele in escaped clones than in clones without LOH (Figure 3D). These results confirm that negative selection for neoantigens shapes the neoantigen landscape of individual subclones inside a tumour.

### Evolutionary dynamics under immunotherapy

We extended our simulations to include immunotherapy after tumour detection, to study how harbouring different evasion mechanisms can influence the efficiency of therapy. The most commonly used agents in immunotherapy target and inhibit immune checkpoint pathways, helping the immune system to overcome immune evasion achieved by PD– L1 or CTLA– 4 overexpression and re– activate immune predation of (neo)antigenic cancer cells. This treatment method had highly promising results in hyper– mutated tumours, which often over–express immune checkpoint proteins^16,38^ – however, some patients do not respond to therapy, or relapse, and the mechanism of resistance is poorly understood. A potential explanation is the presence of an additional immune escape mechanism, such as loss of function alterations in the antigen presentation machinery. We therefore used our model to simulate hyper– mutated tumours with active immune evasion that also harboured a subclone with defective presentation.

We introduced two different types of escape (PDL1 expression and loss of HLA) stochastically during tumour growth (Methods). After the tumour population grew up to detection, we simulated immunotherapy (anti– PDL1) by cancelling the effect of immune– evasion, and also increasing selection pressure to model re– activation of the immune system. Under this scenario, the cell population whose survival relied on active escape (PDL1 expression) rapidly declined – however, small clones that harboured passive– type immune escape (HLA– LOH) continued growing and eventually overtook the tumour (Figure 3E). Neoantigens were progressively pruned from the expanding passively– escaped clone, leading eventually to an immune cold tumour. The same behaviour was observed if a mainly antigen– hot tumour contained a small antigen– cold subclone that underwent sufficient immuno– editing prior to therapy. As a result, any residual tumour post– therapy was predicted to have an immune phenotype distinct from the original tumour, which would likely be resistant to reapplication of the same mode of therapy. Therefore, the modelling predicts that the re– introduction of strong selection through immunotherapy promotes the emergence of resistance, and, combination or evolutionary therapeutic approaches^39^ might be needed to successfully control a tumour in the long term.

### Negative selection leads to effectively neutral evolutionary dynamics

Finally, we sought to explore how negative selection shapes the distribution of subclone sizes within a cancer. We have previously shown that the distribution of variant allele frequencies (VAFs) of somatic mutations provides a way to infer the evolutionary dynamics of individual tumours^7,22^. Specifically, here we aimed to establish the ‘signature of immuno– editing’ on the VAF distribution, as measured in a single individual. We returned to the model of tumour growth to simulate tumours with negatively selected subclones, and analysed the resulting synthetic VAF distributions.

First, we considered the VAF distribution in antigen– hot and – cold tumours separately (Figure 4A– B). We have previously shown that neutral subclone evolution leads to a characteristic distribution of subclonal mutations, whereby the number of mutations at frequency *f* (*M*(*f*)) is proportional to 1/*f*^2^ (or 1/*f* in the cumulative distribution)^7^. In negatively selected cancers, subclonal neoantigens are depleted such that they are rarely present in large subclones, and hence the vast majority of higher frequency mutations in the VAF distribution are neutral passenger mutations, that naturally evolve according to *neutral dynamics* (Figure 4A). In antigen– hot tumours, the same phenomenon is exacerbated: since all tumours cells already carry a negative– selected neoantigen, subclones that then acquire additional neoantigens experience even stronger selection, leading to rapid extinction via immuno– editing.

**Figure 4:**
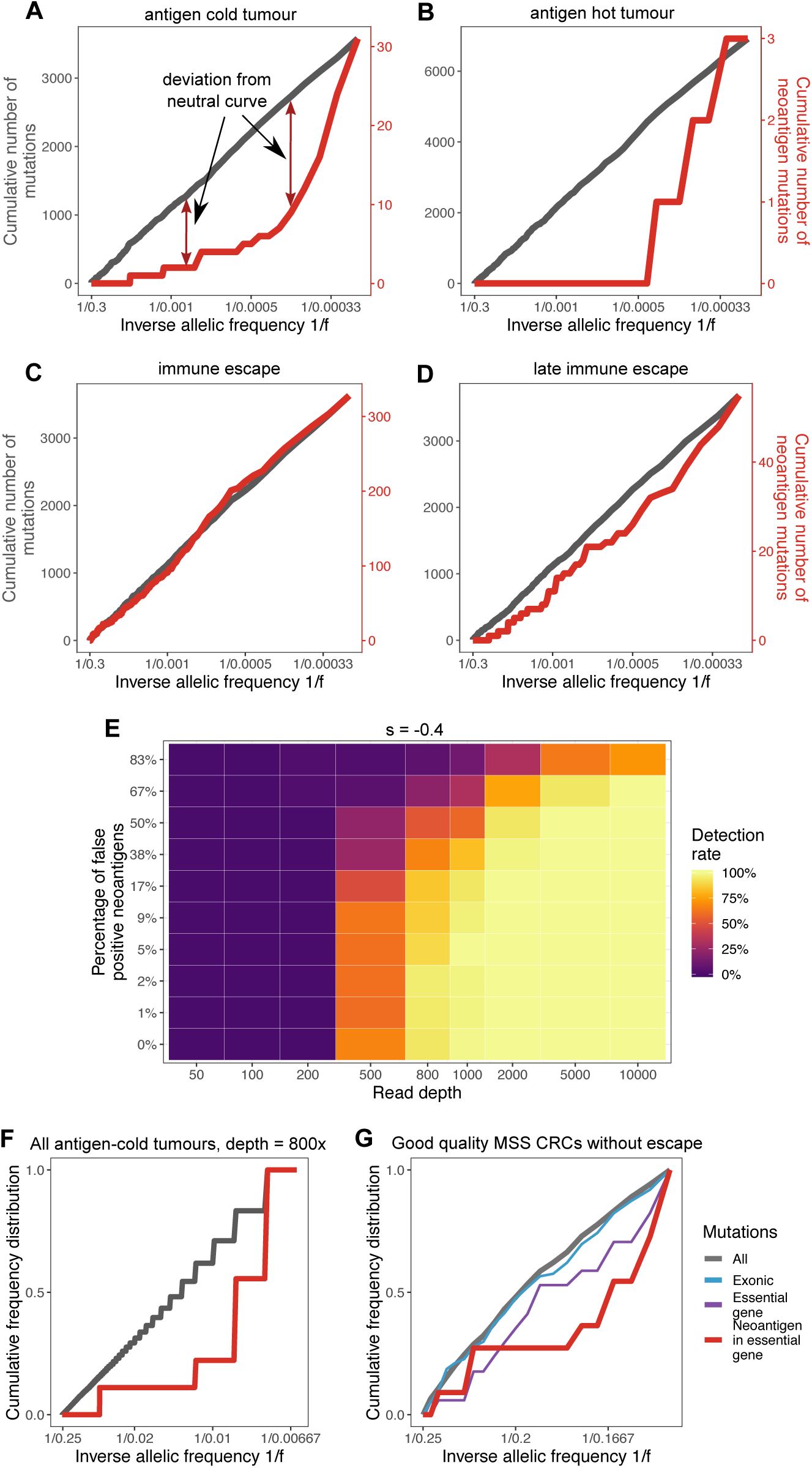
Negative selection leads to effectively-neutral clone size distributions. **(A-D)** Cumulative number of mutations plotted as a function of the inverse of the frequency of all mutations (grey, left y axis) and neoantigen-associated mutations (red, right y axis) harboured in at least 30 cells in (A) an antigen-cold tumour; (B) an antigen-hot tumour; (C) a tumour with clonal immune evasion; and (D) a tumour with subclonal immune escape, introduced at the tumour size of 2000 cells. All tumours are simulated with selection strength s=-0.4. **(E)** Detection rate to identify negative selection from the overall VAF distribution as a function of sequencing read depth (x axis) and false neoantigen rate (y axis). Detection rate is computed as the proportion of 100 simulated tumours with significant difference (Kolmogorov-Smirnov test, α = 0.1) between the distribution of all mutations and neoantigen-associated mutations. **(F)** Synthetic cumulative VAF distribution as a function of the inverse of the frequency of all (in grey) and neoantigen-associated mutations (in red) detected with a sequencing depth of 800x in all antigen cold tumours from a simulated set of 100. **(G)** Cumulative VAF distribution of mutations detected in the filtered subset of TCGA MSS CRCs. The distribution is shown for all mutations (grey), only mutations located in exons (blue), exonic mutations in essential genes (purple) and neoantigen-associated mutations in essential genes (red).

The evolutionary dynamics under negative selection are the inverse of those under positive selection. Under positive selection, the expanding clone carries passenger mutations within the clone to higher frequency than expected from the neutral expectation, and hence positive selection is evident from these passenger mutations at high frequency^22^. Under negative selection, a contracting clone never has the opportunity to acquire passenger mutations, and so the VAF distribution consists of only neutrally evolving mutations. Correspondingly, in both antigen– cold and antigen– hot tumours the VAF distribution for all mutations (grey lines in Figure 4A– B) is dominated by neutral non– antigenic mutations and is not discernibly different from the neutral expectation; and paradoxically the stronger the strength of negative selection, the more neutral– like the evolutionary dynamics.

The VAF distribution computed of solely neoantigens shows depletion relative to the neutral expectation (red lines in Figure 4A– B), consistent with population genetics theory^23,24^ and so in theory negative selection could be detected by considering the VAF distribution of neoantigens. However, we reiterate that negative selection means very few neoantigens persist in the tumour and most of the antigens that persist are at very low VAF. For instance, in the tumour in Figure 4B, neoantigens make up <0.05% of the total detectable mutations (∼3/7000 mutations), despite 10% of all new mutations in the simulation being antigenic. In practice, this means that detecting negative selection from the VAF distribution of neoantigens in tumour sequencing data is problematic, since (a) there are expected to be too few subclonal neoantigens to resolve the distribution, and (b) most antigens are at low VAF where the power to detect variants and accuracy of VAF measurement are both lowest. Moreover, neoantigen identification from DNA sequencing alone has a high rate of false positive calls^40^, and so the VAF distribution of neoantigens is expected to be ‘contaminated’ with a large proportion of neutrally– evolving passenger mutations.

Simulations of clonal immune escape show a curve consistent with neutral dynamics, as is expected (Figure 4C). Subclonal immune escape, experienced when immune escape occurs at a late stage of cancer growth, on the other hand leads to non– neutral, but apparently less selected dynamics. This is due to the escaped subclone effectively experiencing decreased death rate relative to other antigenic tumour cells, and dampening the effect of depletion measured in the total set of neoantigens (Figure 4D).

We performed power calculations on simulated tumours to determine the detectability of negative selection in currently available sequencing data. Real sequencing data naturally introduces uncertainty about mutation VAF due to limited sequencing depth and several sources of sampling bias^41^ and imperfect prediction of antigenicity^40^. We explored the effect of read depth and neoantigen mislabelling on the ability to identify negative selection from the VAF distribution in individual tumours. We compared the distribution of all detected mutations to that of the neoantigen– associated subset using the Kolmogorov– Smirnov test, and identified any samples as under selection in which the p– value of the test was below 0.1. The simulations predicted that very high depth sequencing was required to robustly call negative selection from VAF distributions (Figure 4E). A major pitfall in tumours with high selection was that they contained too few neoantigens to construct the VAF distribution, whilst low selection pressure decreased the confidence in detecting a signal of systemic depletion (Figure S4). Erroneously labelling neoantigens also had a major impact on the detection of negative selection, though this could be mitigated by very high– depth sequencing depth.

In order to overcome the technical issues of limited sequence depth and low antigen numbers, we pooled mutations from groups of tumours and considered their combined CCF distribution. Within the pool, there were adequate subclonal neoantigens to resolve the clone size distribution and detect negative selection (Figure 4F), in a similar manner how cohort– wide positive selection by dn/ds analysis is evaluated^42^. However, the efficacy of pooling strongly relies on the similarity of the tumours included in the cohort, and a heterogeneous mixture of mutations could hinder any selection signal. To this end, we filtered TCGA CRCs to retrieve only MSS samples with tumour content above 75% ^43^, without detected immune escape; and pooled mutation from this cohort. We investigated a specific subset of genes identified as essential genes^44^ that we expect to be both expressed in the cohort and under more controlled selection. The emerging CCF distribution in the subclonal regime, shown in Figure 4G, revealed, as expected, a slight depletion of mutations that are located in essential genes, and a stronger signal in antigenic mutations that occurred in these genes.

### Proportional neoantigen burden can measure selection pressure

As a practical limitation of detecting negative selection lies in the low number of neoantigens, we next investigated if the degree of depletion holds information on the evolutionary dynamics. To study the effect of selection pressure, we needed a measure that could isolate between variations of immunogenic burden arising by chance (e.g. due to different overall mutation burden^45^) and as a result of evolutionary dynamics. Therefore, we calculated the proportional neoantigen burden (the proportion of missense mutations in the tumour that are immunogenic) as a tool to quantify the depletion of neo– epitopes. This quantity is independent of the overall mutation rate or turnover in the tumour population, and only influenced by two rates: (i) the emergence of new neoantigens and (ii) the death of cells carrying neoantigens.

In simulations, we set the proportional neoantigen production rate to a known value *p*_*na*_=0.1 and considered how the negative selection strength (manifested by an increased death rate of antigenic clones) determined the proportional neoantigen burden. Proportional neoantigen burden decreased as the strength of negative selection increased, and as expected, the proportional neoantigen burden mirrored the input ratio in the absence of negative selection (Figure 5A).

**Figure 5:**
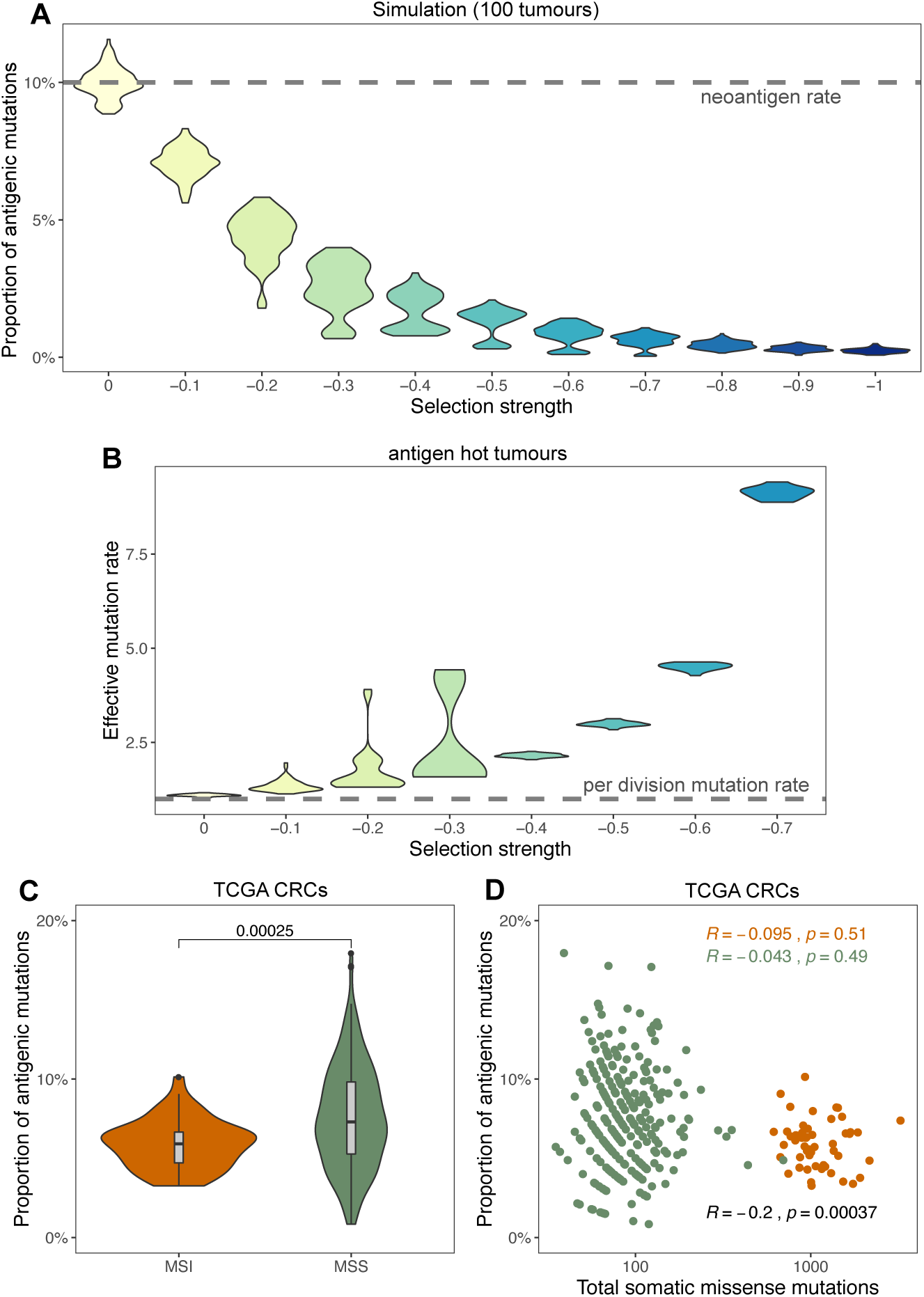
Proportional neoantigen burden as a measure of selection. **(A)** The proportion of neoantigen-associated mutations (the percentage of all mutations) as a function of negative selection pressure, computed from 100 tumours each. The neoantigen generation rate per mutation is indicated with a horizontal dashed line. **(B)** Effective mutation rate (per cell division mutation rate divided by per cell division death rate) computed from the VAF distribution of mutations in antigen hot tumours as a function of negative selection pressure. **(C)** Proportional neoantigen burden of MSS and MSI CRC samples from TCGA. The p-value of Mann-Whitney test is reported above the violin plots. **(D)** Scatter plot of total somatic missense mutation burden and the proportion of neoantigen-associated mutations for the same samples. Results of Pearson’s correlation test are reported for MSS (green), MSI (orange) and all samples combined (black).

We have previously shown that in the case of effectively– neutral subclonal dynamics, the effective mutation rate (µ/ß), defined as the ratio of the per– cell division mutation rate (µ) divided by the per– cell division death rate (ß), is specified by the steepness of the 1/f^2^ VAF distribution^7^. Since negative selection leads to effectively– neutral dynamics, this method can be validly applied to tumours experiencing negative selection. We measured the effective mutation rate as a function of increasing negative selection for neoantigens. Stronger negative selection caused higher effective mutation rates in antigen– hot tumours (Figure 5B). This is because stronger negative selection increased the death rate (ß) of all cells in clonally antigenic populations, increasing the ratio (µ/ß). In other words, stronger negative selection acting on antigen carrier cells leads to more cell division and mutations, between each tumour population size doubling. The apparently high mutation rates observed in hyper– mutated tumours might be a consequence of the same mechanism: a moderate mutation rate could appear to be much higher because of a significantly increased rate of cell death (due to deleterious mutations or immune predation).

We tested the proportional neoantigen burden in TCGA cancers stratified by MSI status (samples with less than 30 missense mutations and with polymerase– ε (POLE) mutations were omitted from the analysis). Relative neoantigen burden was significantly lower in MSI than in MSS samples (Figure 5C), despite MSI cancers harbouring a higher number of neoantigens overall. This suggested that MSI cancers experienced stronger negative selection for neoantigens than MSS cancers. This could be a consequence of the high degree of immune infiltration in MSI tumours relative to MSS cancers^46^. The relation could also emerge if there was a negative correlation between overall missense mutation burden and relative neoantigen burden. However, no such negative correlation was observed (Figure 5D) therefore we conclude that via proportional neoantigen burden, we quantified differences in selective pressures between the two groups of tumours.

## DISCUSSION

Here we have investigated the evolutionary dynamics of neoantigens and immune escape in growing tumours using a simple mathematical model of tumour evolution. We validated the model against genetic data characterising the neoantigen burdens in colorectal cancers. Our analysis shows how negative selection by the immune system (immuno– editing) sculpts the clonal architecture of the tumour in a subtle way: the hallmark of negative selection is a lack of neoantigens at intermediate frequency within a tumour, and conversely, that the presence of antigens at intermediate frequency is a hallmark of immune escape. Moreover, strong negative selection for neoantigens effectively inhibits tumour growth, but inevitably provides a strong selective pressure for the evolution of immune escape. Consequently, the observation that many colorectal cancers are both highly (neo)antigenic and also have immune escape points to a critical role for immune evasion in the genesis of malignancy. Further work directly measuring the immune repertoire at the time invasion first occurs is now required.

We showed that negative selection is detectable via relative depletion of neoantigens, giving rise to a characteristic ‘signature’ VAF distribution of the mutations under selection. Our simulations agree well with theoretical predictions for clone size distributions under purifying selection ^23,24^. However, we show that both weak and strong selection (depletion) are hard to detect at the currently available sequencing resolutions. The former due to an evolution following effectively– neutral dynamics, and the latter because strong depletion leads to an insufficient number of disadvantageous mutations. However, recent evidence suggests that mutations present in a low proportion of cells might not elicit an immune response depending on the characteristics of the arising neopeptides^47^, and we expect this effect to be reinforced in tumours with complex spatial architecture such as colorectal cancer which allows cancer cells to ‘hide’ from the immune system. This means that negative selection only operates on larger clones – and we note that potentially this shift from neutral to negatively– selected dynamics could be detected with very high depth sequencing or over a sufficient cohort of cancer cases.

Nevertheless, negative selection may be evident in clone– size distributions at the cohort level (despite not being detectable in individual tumours) wherein pooling data from multiple tumours allows for sufficient mutations to resolve clone size distributions. Applying a pooling method across a cohort of TCGA CRCs assumed to be of similar immune– phenotype, we have detected a trend of negative selection in both essential genes and antigenic mutations harboured in essential genes. This is consistent with the previous observation that binding affinity to the MHC determines which driver mutations will be found in cancer, and that this immune filtering occurs differentially between MHC– I and MHC– II ^48,49^. The approach of testing for relative neoantigen depletion could also be applied to test other sets of mutations presumed to be under negative selection, such as genes encoding essential cell functions, natural antigenic peptides^50^ or modified peptides presented to CD4+ T– cells through the MHC– II complex.

Our simulations show that under negative selection, the overall VAF distribution of a tumour will be effectively– neutral, since only neutral passenger mutations survive immune predation and are able to spread through the tumour. Paradoxically, the clone size distribution becomes more neutral– like as the strength of negative selection increases, as negatively selected clones cannot become established in the tumour and so have no effect in the clone size distribution. Together, these results highlight a practical issue for detecting negative selection in currently available sequencing data: negative selection causes only very slight deviations from the neutral expectation in the overall mutational spectrum, while specifically detecting selection requires correct characterisation of mutations into putative neoantigens and non– antigenic mutations. The latter is still an open problem as many steps of the immune presentation process are not well understood, and therefore current methods might be identifying a high number of false positives, whilst also missing other significant neoantigens, e.g. spliced peptides^40,51^.

Our findings have potential implications for the stratification of patients for immunotherapy. Our simulations show that tumour mutational burden (TMB) may not necessarily be the best predictor of immune– checkpoint blockade response. This is because effective immune surveillance (e.g. no immune escape) leads to the killing of antigen bearing clones, with the net result of increasing the effective mutation rate of the surviving tumour cells, and enrichment for non– antigenic mutations. Thus, we predict that tumours with a high TMB and without evidence of evolved immune escape mechanisms are unlikely to respond to immune blockade therapy. We predict that tumours with clonal neoantigens are very likely to have evolved immune escape, particularly if the patient’s immune system is highly predatory. In such cases, our model predicts the observation that clonal antigens elicit sensitivity to immune checkpoint blockade ^34^. Our modelling also predicts that immune therapies ‘targeted’ against a specific neoantigen (e.g. CAR– T therapies) can only hope to affect a cure if directed against clonal neoantigens, otherwise a subclone without the neoantigen target will experience net positive selection when the therapy is applied^52,53^. Relatedly, a subclone that escapes immune blockade therapy and reforms a tumour is predicted to have a very different immuno– genotype/phenotype to the original tumour due to the action of immune predation during clone emergence, with ramifications for metastatic potential and choice of second– line therapy.

In summary, we have provided a mathematical framework to explore the evolution of neoantigens in growing tumours, and illustrated how quantitative understanding of tumour– immune co– evolution can provide new insight into the choice and effectiveness of immunotherapies.

## METHODS

### Mathematical model of tumour growth and mutation accumulation

We created a minimal model to capture the growth of a tumour population and accumulation of mutations under selection pressure from the environment. Since the scope of the model was to match information derived from bulk sequencing of tumours, we chose to only model explicitly the proliferation, death and mutation of tumour cells, and include environmental information (e.g. the level of T– cell infiltration) implicitly through appropriately chosen model parameters. In order to capture the inherent randomness in the modelled events, we used a stochastic birth– death process. To reduce computational burden, we applied a rejection– kinetic Monte Carlo algorithm ^22,54^ that allowed for the efficient simulation of large populations of cells with identical birth/death kinetics. The steps of the simulation algorithm are detailed below and illustrated in Figure 1A.

First, a single progenitor cell is defined that already carries a set of mutations providing it with sufficient growth/survival advantage to outgrow a normal cell population. Each of the cell’s mutations have a unique identifier, and that cell has an intrinsic immunogenicity value determined by its mutations. Starting from this single– cell tumour, in each simulation step a cell in the population is selected, and that cell undergoes one of three possible life events:

– *Proliferation*: The cell divides and gives birth to two daughter cells. These cells carry all mutations and information contained in the mother cell, but also acquire new mutations. For each newly generated mutation, it is randomly decided whether the mutation is antigenic.

– *Death*: The cell dies and is removed from the population.

– *Waiting*: No proliferation or death event happens; the cell is not altered in any way.

The probability of each event is defined by the cell’s proliferation and death rate (*b* and *d*_*i*_) as *b*/(*b* + *d_max_*), *d*_*i*_/(*b* + *d*_*max*_) and 1 – (*b*+*d*_*i*_)/(*b* + *d_max_*), respectively.

In proliferation events, each daughter cell gains *N*_*m*_ new, independent mutations, where *N*_*m*_ is sampled from a Poisson distribution with parameter *µ*, the cell’s mutation rate. Antigenicity is randomly assigned to newly generated mutations according to the antigen production rate, *p*_*na*_; the probability that a newly generated mutation has immunogenic properties.

The above step of randomly selecting a cell and one of the three possible events is repeated until the tumour reaches a predefined population size (representing the tumour reaching a clinically detectable size, for simplification we set it to 105 cells) or sufficiently long time elapsed without tumour establishment (corresponding to no cancer formation in the patient’s lifetime, set to 300 time units).

For simplicity, we assumed that each cell had the same proliferation rate, *b* = 1, and cells with different properties in the tumour could only differ in their death rate determined by the immunogenicity status of the cell. As the average dynamics of a subclone’s growth are determined by the ratio of its birth and death rate, this simplification does not compromise the generality of the results derived from the model. Cells were considered immunogenic if they carried at least one (neo)antigenic mutation. We modelled the effect of immune system on neoantigen– carrier cells as an increase in cell death probability, representing additional T– cell mediated death. Non– immunogenic cells were assigned a basal death rate of d_b_ = 0.1, to account for any detrimental mutations present in the first progenitor cell and immune– independent cell death. Immunogenic cells, on the other hand, had an increased death rate, *d*_*i*_, computed as *d*_*i*_ = (1 + *s* * *n*_*i*_) *d*_*i*_ – 1 + 1. Note, that *d*_*i*_ represents a summary of many processes not modelled explicitly, such as (i) sufficient presentation of neoantigens on the cell surface; (ii) recognition of neoantigens by T– cell; (iii) active killing of carrier cells initiated by recognition. As each of these processes have inherent randomness (e.g. the chance meeting of a neoantigen– carrier and an approriate T– cell is necessary for recognition), similar selection pressures can arise from different combinations of the above. We chose to integrate all variability into a single probabilistic rate to be able to observe general qualities of the tumour– immune interaction without the need for precise parametrisation.

In an extension of the model, we also considered the acquisition of immune escape through tumour growth. Immune escape was modelled as a heritable property of a cell, gained through either escape– associated mutations that were sampled from newly generated mutations, similarly to neoantigens, with probability *p*_*esc*_; or through manual introduction of the escape alteration at a pre– determined clone size to achieve clonal or subclonal immune escape. We considered two different types of escape mechanism: (i) active evasion, which shields the cell from negative selection (decreasing its death probability to *d_b_*) but leaves neoantigen– carrier cells highly immunogenic (corresponding to escape mechanisms such as PD– L1 overexpression); and (ii) passive evasion, which lowers the cell’s immunogenicity in the presence of neoantigens (representing escape mechanisms arising from loss/mutation of antigen– presentation machinery).

We chose this level of abstraction in the model to enable us to investigate evolutionary paradigms on a general level, without having to rely on precise parameterisation of many sub– processes. Therefore, while the reactions and parameters included in the model might not correspond to a single biological event, the model can provide a qualitative description of a high range of tumour– immune environments with appropriate choice of parameters. Furthermore, the model can be easily extended to account for further biological processes, in the same manner in which immune escape was included, to provide predictions on therapeutic interventions.

### Choice of model parameters

We chose model parameters to represent a wide range of possible tumour– immune environments, and be easily interpreted in terms of growth dynamics. Therefore, all rate values time units in the simulation are scaled to the rate of proliferation/birth events (which is 1), and for example the death rate of 0.9 or 1.2 correspond to overall population growth or decline, respectively.

The probability of proliferation, death and waiting events (see steps above and Figure 1A) for a cell with birth rate b and death rate *d*_*i*_ were *b*/(*b* + *d_max_*), *d*_*i*_/(*b* + *d_max_*) and 1 – (*b*+*d*_*i*_)/(*b* + *d_max_*), respectively. Due to the linear connection between the number of neoantigen harboured by a cell, and its death rate, *d_max_* was computed from the maximum number of simultaneous antigens in any cell in the population.

Mutation rate values (*µ* and *p*_*na*_) were fixed for the entire population in the beginning of each simulation. To correspond to tumour samples that were sequenced using whole– exome sequencing, we set the mutation rate relatively low, *µ* = 1, with the exception of hyper– mutated cancer, which had a higher mutation rate, *µ* = 6. This effectively meant that not all cell divisions introduced new (exonic) mutation in the daughter cells. The value of neoantigen probability, *p*_*na*_, by default was set to 0.1 based on the experimental observation that roughly 10% of mutations that induce a change in protein– sequence resulted in new peptides identified as neoantigens. However, we explored values of 0.2 and 0.05 as well, and found model predictions were in good agreement with those derived using the 10% setting.

Alterations that induce immune escape are rare, because these can only arise from a very limited set of exonic loci. We therefore set *p*_*esc*_ to 1×10^−1^ for all immune escape alterations. The decrease in immunogenicity in passive evasion was set to 0.9. This decrease in antigenicity indirectly affected death, as we assumed that only a fraction of the total number of antigens harboured in the cell remained antigenic, and updated the number of antigens, *n*_*i*_, accordingly.

In summary, the following parameters were used in all simulations: *b* = 1; *d*_*b*_ = 0.1; *µ* = 1 (not hyper– mutated) and *µ* = 6 (hyper– mutated); – 1 ≤ *s* ≤ 0 (as indicated in figures or in caption); *p*_*na*_ = *0.1* (high) and *p*_*na*_ = *0.05* (low antigen rate); *pesc* = 1 × 10^−6^ (where applicable).

### Simulation of CCF values

At the end of each tumour growth simulation, mutations harboured in currently living cells (identified by unique mutation ids) were collected in a dictionary with their respective abundance: the total number of cells harbouring the mutation. Only mutations harboured in at least 10 cells out of 10^5^ were considered in any analysis. CCF values were either computed by taking the raw frequency values (mutation abundance divided by the total tumour population size), or via a simulated sequencing step introducing noise to these frequencies with indicated read depth. For a given read depth, *D*, each frequency value, *f*, was substituted by a new frequency sampled from a binomial distribution with parameters *D* and 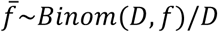. Many mutations will have a sequenced frequency of 0, corresponding to mutations that are not picked up due to limited detection power. Therefore, we filtered for mutations with 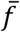 above 0, effectively enforcing a simulating a depth– based detection limit.

### TCGA sample acquisition and processing

All samples from the TCGA COAD and READ (merged together as CRC) were retrieved through the NCI Genomics Data Commons (GDC) portal^55^. Only patients with available matched germline information (from blood samples) were considered. For each sample, purity (fraction of tumour cells in the sample) and overall ploidy were evaluated using ASCAT^43^ on Affymetrix SNP array data. Samples with purity below 0.4 and ploidy above 3.6 were excluded from further analysis, leaving 363 CRC samples for which HLA typing and neoantigen calls were performed (Table S1). For analysing immune escape, the cohort was narrowed down to patients for whom gene expression data was available in GDC; and at least one pair of their HLA A/B/C alleles were heterozygous and distinct enough to allow for loss of heterozygosity calls (n = 339).

For each patient considered, the following information was downloaded: blood derived normal bam files; primary tumour bam files; variant call (vcf) files processed with Mutect2; SNP array files; gene expression HTSeq counts (where available); and clinical information. We used the controlled– access variant call format (vcf) files to avoid over–filtering and missing antigenic variants. The variants were filtered to only include variants not present (allelic depth of 0) in normal samples.

Samples were divided into MSS, MSI and POLE subtypes using data integrated from clinical TCGA annotation^56^, calls derived for the software MANTIS^57^, and mutational signature activities computed using non– negative least squares regression^58,59^. Samples with a MANTIS score ≥ 0.5 and TCGA annotation of ’MSI– H’ (where available) were considered MSI, and those with MANTIS < 0.5 and ‘MSI– L’/‘MSS’ were labelled MSS. In case the two sources of information contradicted each other, neither of the categories was assigned. Samples with at least 1500 mutations inferred to originate from the characteristic POLE signature (signature 10 in Martincorena et al.^58^) labelled as POLE tumours regardless of their MSI status.

### Multi–region sequenced dataset processing

The multi– region sequenced colorectal dataset was accessed from Cross et al.^37^. Raw data is available from European Genome– Phenome Archive (https://ega–archive.org/) at accession code: EGAS00001003066. Bam files with marked duplicates were used for HLA calling and HLA variant detection. As in the original work, variants were called using Platypus^60^, annotated by ANNOVAR^61^, and filtered to only contain somatic single nucleotide variations that were present in at least 1 tumour sample and in either 0 reads in the normal sample (for normal coverage <=40 reads) or in at most 1 read (for normal coverage above 40 reads).

### Computation of VAF and CCF

For each mutation, we calculated the variant allele frequency (VAF) as the number of mutant reads spanning the position, divided by the number of total reads of the position. The proportion of cancer cells carrying a particular mutation (cancer cell fraction, CCF) was calculated from the VAF of the mutation, sample purity (tumour content), and copy number (CN) of the mutation’s genomic locus as: (*VAF* * *CN*) purity. CCF values above 1 (arising from sequencing noise and copy– neutral loss– of– heterozygosity events) were assumed to be 1.

### HLA haplotyping and calling immune escape

HLA– A, – B and – C haplotyping was performed on blood derived normal bam files using POLYSOLVER, which performs high precision HLA– haplotyping on whole– exome sequencing data, and also enables subsequent mutation detection based on the inferred alleles for detection of mutations in the highly polymorphic HLA genes^30^. As POLYSOLVER takes into account the patient’s race to compute the likelihood of each allele haplotype, we supplied ethnicity data, where available from clinical TCGA information, and ran haplotyping with race ’Unknown’ otherwise. Mutations in HLA alleles were called using the specialised variant calling and annotation functionality of the same software, using the previously called HLA haplotypes. Variant calling was run using default settings and HLA was considered mutated if at least one allele had a nonsynonymous somatic mutation located in an exon or at a splice– site. Mutations in B2M were called if the sample contained a nonsynonymous somatic mutation located inside one of the exons of the B2M gene, as annotated by ANNOVAR^61^ and confirmed using Variant Effect Predictor^62^.

Loss of heterozygosity at the HLA locus was assessed using the software LOHHLA, which permits allele– specific copy number estimation using reference information specific to the particular HLA haplotype^28^. Blood derived normal, and tumour bam files were used. Tumour purity and ploidy estimates were derived from ASCAT (for TCGA data) and from Sequenza (for the multi– region sequenced colorectal tumours). A sample was considered to have Allelic Imbalance at an HLA locus if the corresponding p– value was below 0.01 and LOH if, in addition, the copy number prediction of that allele was below 0.5, with the confidence interval strictly below 0.7.

Immune checkpoint over– expression was assessed using RNA– seq data. Normal expression values (in transcripts per million (TPM)) of PD– L1 and CTLA– 4 were established for each cohort from TCGA based on RNA– seq counts of the two proteins in ‘solid tissue normal’ samples. Checkpoint over– expression was called if either PD– L1 or CTLA– 4 expression in the tumour was higher than the mean plus two standard deviations of normal expression.

### Neoantigen prediction

Neoantigens were predicted from variant call tables and HLA types using NeoPredPipe^31^, a neoantigen prediction and evaluation pipeline designed for parallel analysis of single– and multi– region samples. The pipeline was run with default analysis settings and preserving intermediate files (–p flag), using hg38 and hg19 ANNOVAR^61^ reference files for annotation of the TCGA and multi– region CRC samples, respectively. The analysis outputted a table of novel peptides binding the patient’s MHC– I molecules and their respective recognition potential calculated from their MHC– binding affinity and similarity to pathogenic peptides, as described in ^32^. Unless stated otherwise, we labelled a peptide as neoantigen if its recognition potential was >= 10^−1^ to focus on antigens with the highest predicted probability of eliciting an immune response: both similar to known pathogens and significantly stronger MHC– binders than their wild– type counterpart. Similarly, a mutation was considered (neo)antigenic if at least one altered peptide produced from the mutated genomic region was a neoantigen. Essential genes, and neoantigens located in essential genes were identified using the list of shared genes in ^44^.

### Statistical analysis

All data processing and statistical tests were performed in R (version 3.5.0) using built– in functions. The tests and functions used were as follows: Figure 2C: Mann– Whitney U– test/ Wilcoxon sum– rank test (*wilcox.test*, default settings). Figures 2D and S3: Chi– squared test (*chisq.test*). Figure 2E: One– sided Mann– Whitney U– test (*wilcox.test* with option *alternative=’greater’*). Figure 3C and D: One– sided Wilcoxon signed– rank test (*wilcox.test with options paired=TRUE and alternative=’greater’*). Figure 5E: Kolmogorov– Smirnov test (ks.test) between the raw VAF distribution of neoantigens and all mutations. The two distributions were deemed significant if the p– value was below 0.1. Figure 5C: Mann– Whitney U– test/ Wilcoxon sum– rank test (*wilcox.test, default settings*); Figure 5D: Pearson’s correlation test (*cor.test*).

On all boxplots presented (Figures 2E, 3C and 5C), visual elements correspond to the following summary statistics: centre line, median; box limits, upper and lower quartiles; whiskers, 1.5x inter– quartile range; additional points, outliers below/above 1.5x inter– quartile range.

## ACKNOWLEDGEMENTS

This work was supported by the Wellcome Trust (202778/B/16/Z to A.S.; 202778/Z/16/Z to T.A.G.; 105104/Z/14/Z to the Centre for Evolution and Cancer, Institute of Cancer Research; 108861/7/15/7 to R.O.S.; 097319/Z/11/Z to C.P.B.) and Cancer Research UK (A22909 to A.S.; A19771 to T.A.G. supporting E.L.). A.S. also acknowledges support from the Institute of Cancer Research (Chris Rokos Fellowship in Evolution and Cancer). A.R.A.A. and C.G. received support from the US National Institutes of Health National Cancer Institute (grant no. U54CA143970). R.O.S. was also supported by the Wellcome Centre for Human Genetics (grant no. 203141/7/16/7). B.W. was supported by the Geoffrey W. Lewis Postdoctoral Training fellowship.

## AUTHOR CONTRIBUTIONS

E.L., A.R.A.A., A.S. and T.A.G. conceptualised the study. A.R.A.A., A.S and T.A.G. acquired funding for the project. E.L., and T.A.G. lead the investigation and wrote the original manuscript. E.L., M.J.W., W.C.H.C., B.W., R.O.S., C.G., J.H., M.R.T., C.P.B. and T.A.G. contributed to the mathematical model, computational framework and bioinformatics analysis. All authors reviewed and approved the final manuscript.

## DATA AVAILABILITY

The datasets analysed during the current study are available from the NCI Genomics Data Commons Portal (https://portal.gdc.cancer.gov) COAD and READ domains, and from the European Genome– Phenome Archive (https://ega–archive.org/) at accession code: EGAS00001003066.

## CODE AVAILABILITY

Julia (https://julialang.org/, version 0.5+) code implementing simulations of the tumour growth model is available from GitHub (https://github.com/elakatos/CloneGrowthSimulation).

## CORRESPONDENCE

Further information and requests for resources should be directed to and will be fulfilled by corresponding author Trevor Graham.

