## Supplementary Figures for "Evolutionary dynamics of neoantigens in growing tumours"

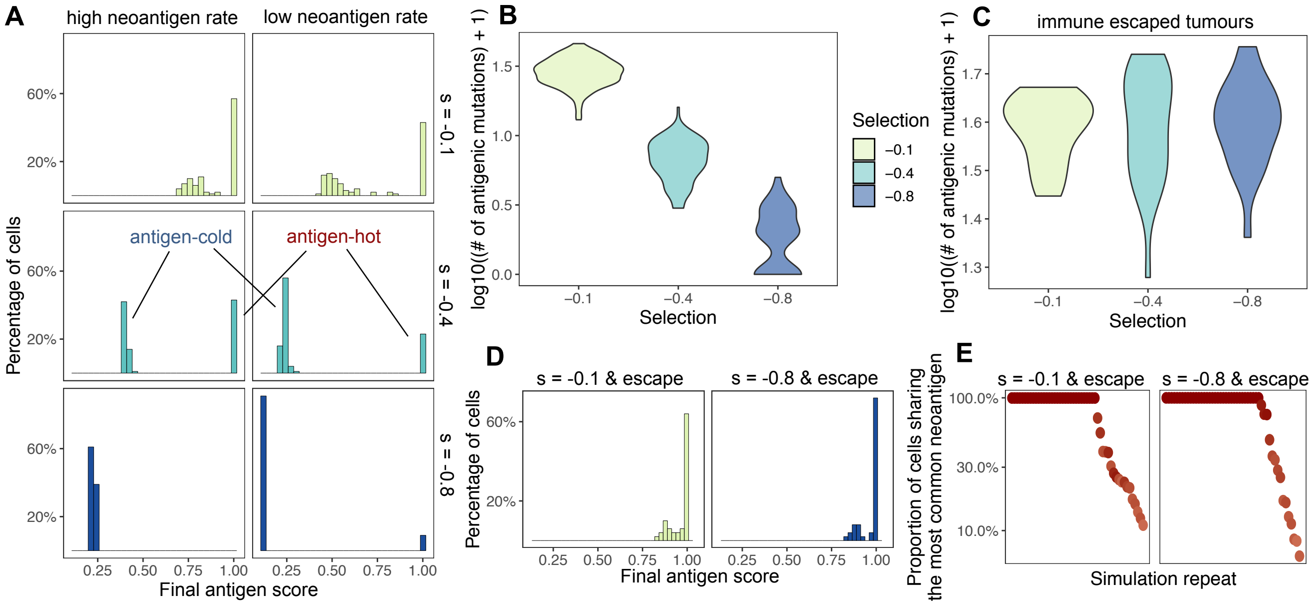


**Supplementary Figure 1: Population-level statistics under varying levels of selection and immune escape. (A)** Distribution of antigen scores of tumour cell populations when reaching $10^5$ cells, for different selection strengths and neoantigen generation probability. 100 tumours were simulated for each parameter combination. **(B)** Distribution of the number of detectable neoantigen-associated mutations (at simulated sequencing depth of ~50x) in 100 simulated tumours each under low, medium and high selection pressure (s = -0.1 (yellow), s = -0.4 (teal), s= -0.8 (blue), respectively). **(C)** Distribution of the number of detectable antigenic mutations in tumours with a clonal immune escape alteration, under three different selection pressures, as in panel B. **(D)** Distribution of final antigenicity values in simulated immune escaped tumours at selections s=-0.1 (low) and s=-0.8 (high). **(E)** Frequency of the most shared antigenic mutation in 100 immune escaped tumours at low and high selection pressures.

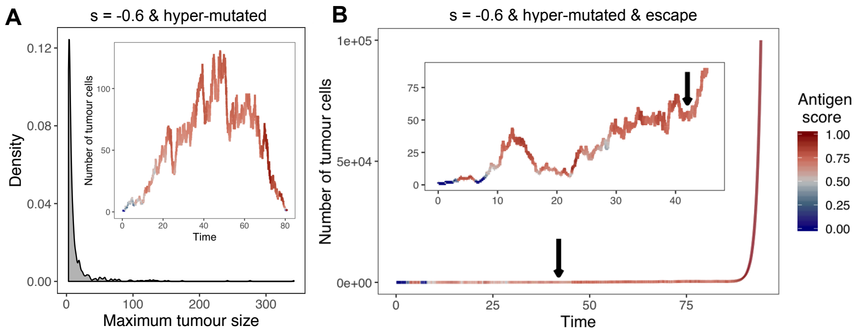


**Supplementary Figure 2: Hyper-mutated tumours are highly antigenic and rescued by immune escape alterations. (A)** Distribution of the maximum tumour size (measured as the highest number of tumour cells) reached by simulated hyper-mutated tumours under selection s = -0.6. The inset shows the growth curve of an example hyper-mutated tumour. The colour of the graph at a given time-point shows the antigenicity of the tumour population (see panel B for legend). **(B)** Growth curve of a hyper-mutated tumour under high selection that acquires immune escape at time 42 (indicated by black arrow). The inset shows the first segment of growth.

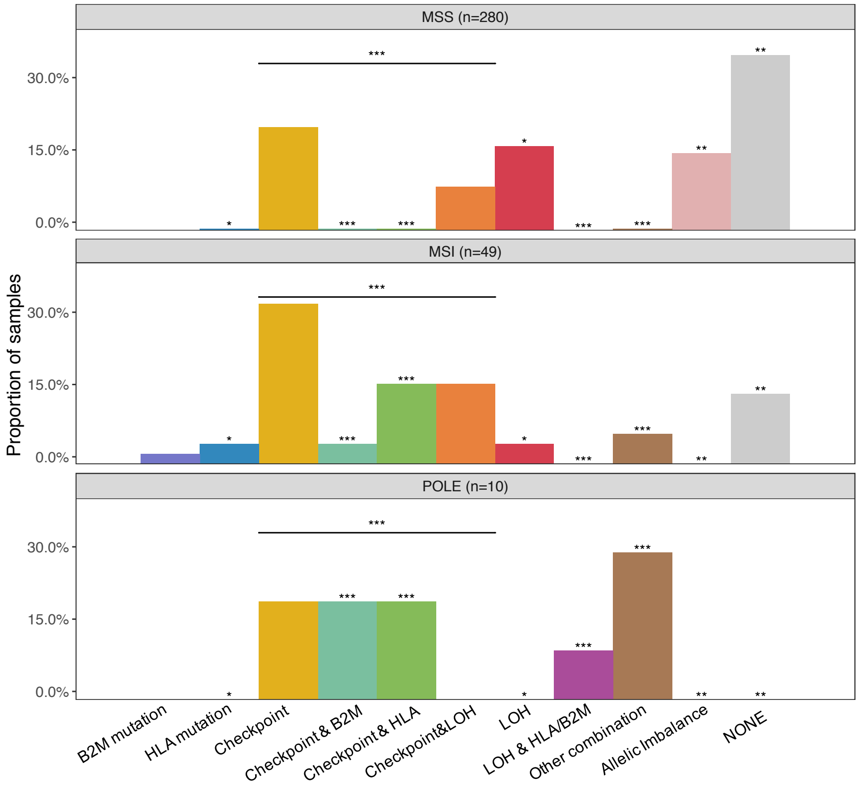


**Supplementary Figure 3: Immune escape in TCGA colorectal cancers.** Prevalence of the individual escape mechanisms considered. Significance markers on top of each bar indicate the result of chi-squared test for that mechanism. An additional test comparing the presence/absence of any immune checkpoint escape is also indicated above the checkpoint columns. (*: p < 0.05, **: p < 0.01, ***: p < 0.001)

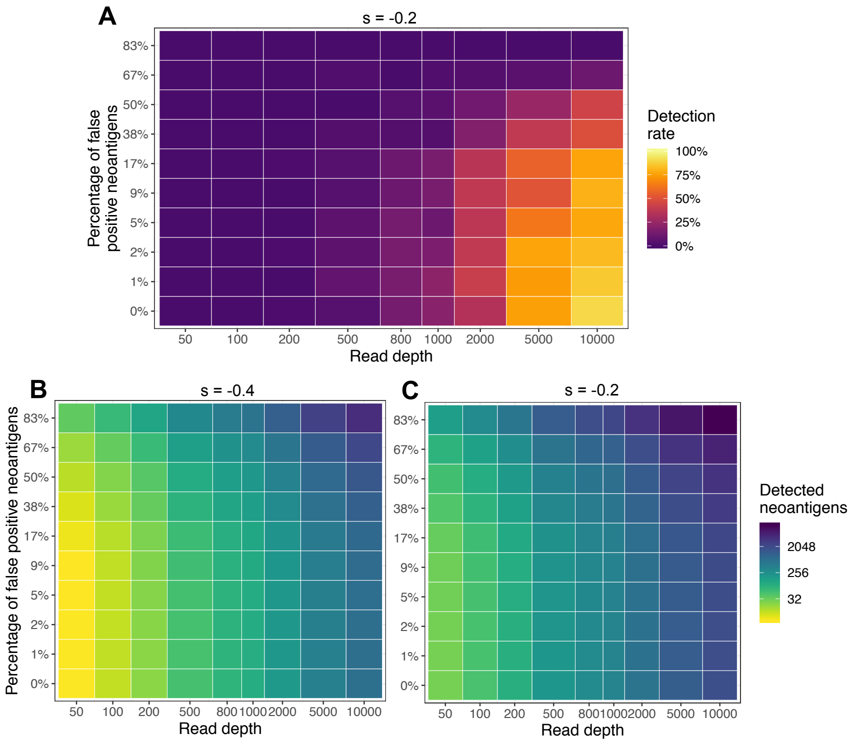


**Supplementary Figure 4: Limits of detection of negative selection for different selection strengths. (A)** Detection rate to identify negative selection depending on sequencing read depth (x axis) and false neoantigen rate (y axis), the percentage of mutations labelled as neoantigens that are false positives, at selection strength s = -0.2. Detection rate is computed as the proportion of 100 simulated tumours with significantly different neoantigen distribution, compared to the total set of mutations (Kolmogorov-Smirnov test, α = 0.1). **(B-C)** Mean number of detected neoantigens (including true and mislabelled neoantigens, and on a logarithmic scale) at each read depth and false neoantigen rate. Sequencing depths below 500x can only detect a few tens of mutations, insufficient for establishing the VAF distribution.

**Supplementary Table 1 (separate file): Colorectal cancer samples in The Cancer Genome Atlas included in the bioinformatic analysis.** The following information is listed for each sample in a tabulator separated format: patient identifier, cancer type (colon or rectal adenocarcinoma), subtype (microsatellite stable (MSS), microsatellite unstable (MSI) or polymerase-ε mutated (POLE)), predicted escape type (as shown in Figure S3), number of antigenic mutations, number of somatic missense mutations, average ploidy and tumour purity (as determined using ASCAT). Fields that could not be determined (MSI/MSS status or immune escape mechanism) are denoted by NA.
